# Investigating the impact of edge weight selection on the pig trade network topology

**DOI:** 10.1101/2024.08.12.607545

**Authors:** Gavrila Amadea Puspitarani, Yan-Shin Jackson Liao, Reinhard Fuchs, Amélie Desvars-Larrive

## Abstract

Traceability of animal movements and robust surveillance are crucial for identifying and controlling animal diseases. Risk-based surveillance, i.e. network-based approaches, can identify higher-risk holdings or trades. However, node ranking, useful to identify ”influential” nodes (holdings) in the network, varies with the considered metrics.

We use a dataset of pig movements in Upper Austria from 2021 to study the robustness of node ranking through three centrality metrics and compare them with epidemic model ranking. Incorporating edge weights may influence the network analysis, therefore, we simulate two representations using edge weights based on: i) the frequency of exchanges between holdings (”frequency-based”) and ii) the number of pigs exchanged (”volume-based”). We compare the impact of the edge weight on the network topology, community structure, and node ranking in a network with 5,766 nodes and 92,914 edges. Results revealed distinct edge weight distributions: frequency-based network exhibited a bimodal pattern, while volume-based was more uniform. Strength centrality exhibited the highest correlation with simulation-based rankings, particularly for the top 5% highest-ranked nodes (*τb* = 0.51 for frequency-based and *τb* = 0.5 for volume-based). These findings highlight that using strength centrality to identify critical nodes can significantly enhance surveillance strategies, making them more efficient and field-deployable, enhancing traditional methods without requiring extensive simulations.

**Author summary:** Early detection of infectious diseases through surveillance activities is important to prevent severe impacts on the livestock sector and avoid major economic losses that often result from delayed detection. Prioritizing surveillance efforts through a data-driven approach presents a strategic advantage compared to random sampling. Here, we propose two network representations of the Upper Austrian pig trade network 2021, which show livestock holdings as nodes and animal movements as links between these nodes, which can represent the volume of pigs exchanged or the frequency of these exchanges. Using network analysis methods, we identify ”influential” holdings in the network, i.e., those playing a key role in the network, either due to their numerous connections with other holdings, their position on many trade paths, or their crucial role in disease transmission dynamics. We show that using strength centrality can effectively identify key holdings for targeted surveillance. Adopting network-based surveillance can facilitate resource allocation for veterinary surveillance programs, offering a cost-effective strategy for disease management, enabling tailored, more effective, and timely interventions.

## Introduction

The transmission dynamic of infectious diseases is tightly linked to the patterns of interaction among individuals [1]. While models of epidemic spread on networks typically simplify these interactions by treating them as binary (i.e. either present or absent) [2], this oversimplification of the epidemiological process overlooks the diversity of the interactions. Hence, it fails to recognize that some individuals engage more intensely in social relationships than others, therefore contributing more to the spread of infectious diseases within the network.

The intensity of the interactions, represented as edge weights in the network, can significantly impact epidemic spread [3]. To accurately capture the contagion process on networks and estimate epidemic metrics, it is imperative to move beyond binary consideration within homogeneous-mixing models and account for the weighted nature of the interactions. This involves examining the distribution of the number of contacts between pairs of individuals and the nature of their interactions [4, 5]). For example, when using human sexual data to understand sexually-transmitted disease outbreaks, both the number of partners and the number of sex acts per unit of time (reflecting the intensity of the interaction) vary among partnerships. Consequently, weighting contacts is essential to accurately capture the heterogeneity in sexual activity intensity per individual, with a significant impact on the dynamics of disease spread within the population [6].

Mobility datasets offer the flexibility to assign different types of weights to links in the network, e.g. edge weight may represent the intensity or capacity of the relationships among network entities [7]. Recent studies in veterinary epidemiology have highlighted the importance of incorporating edge weights when analyzing livestock trade networks, e.g., in the USA [8], Germany [9], and North Macedonia [10], as well as in modeling disease transmission in sheep populations [11]. Büttner et al. [3] reported that the type of edge weights significantly impacts the final epidemic size. Within an animal trade network, edge weights can represent either the number of trades (hereafter referred to as ”frequency-based”) or the number of traded animals (”volume-based”) between pairs of holdings. These values can further be aggregated over specific time intervals (e.g., daily, monthly, or the entire observation period).

Understanding the transmission dynamic is essential in veterinary epidemiology. Frequency-dependent disease transmission implies that the contact rate (and therefore the likelihood of transmission of the disease) is independent of the population size or density. In contrast, density-dependent disease transmission occurs when contact rates between individuals increase as the population density rises, leading to a subsequent increase in disease transmission [1, 12, 13]. Vector-borne and sexually transmitted diseases are generally described using frequency-dependent transmission [1, 5, 14].

Directly transmitted diseases (e.g., classical swine fever, pseudorabies), in contrast, are typically expected to spread in a density-dependent manner [1, 15].

A key challenge in network epidemiology is to identify nodes that have the greatest influence on disease propagation. Node ranking methods have primarily focused on technical measures, often neglecting to verify the actual influence of these nodes during an epidemic process. While studies have demonstrated the existence of superspreaders in specific networks, these findings lack generalizability across diverse real-world scenarios [16–18]. There is currently no consensus on the most effective node centrality measure for quantifying epidemic importance; the choice of metric should be tailored to specific networks and epidemiological contexts. Additionally, the network structure significantly impacts disease transmission. For instance, different networks exhibit diverse distributions of the number of contacts (called degree centrality distribution) across nodes; in many real-world networks, most nodes display a relatively small number of connections while few nodes are highly connected [9, 19–21], acting as ”hubs”. In network epidemiology, such central nodes can potentially be used as sentinels for disease surveillance [22, 23].

In this study, we first aimed to explore, from a veterinary epidemiology perspective, whether the selection of edge weights impacts both the topology of an animal trade network and node ranking based on weighted centrality metrics. Our second objective was to determine whether node rankings derived from centrality metrics in each network aligned with the ranking obtained from a simulation-based contagion process, used as a benchmark. To achieve these objectives, we used the daily records of movement data of pigs in Upper Austria in 2021 and methods from social network analysis [22]. We used a weighted directed network representation of this data, where nodes represent pig holdings and edges signify trades. We created two representations of this network that allowed us to compare two weighting approaches. In the first representation, edge weights represented the number of trades between holdings (”frequency-based” network) whereas in the second approach, edge weights represented the number of animals traded (”volume-based” network).

We investigated the federal state of Upper Austria (1.53 million inhabitants, 11 982.67 km2 [24]) because it contributes 41.7% to the total pig production in the country, exhibits the highest density of pig farms in Austria, and its network of pig movements is more densely connected than in the rest of the country [25].

These characteristics create an interesting opportunity for testing surveillance and control strategies that may not be readily applicable to regions with a sparser network. By leveraging a real-world network, our goal was to provide insights that closely mirror real field conditions with practical applications for disease surveillance and prevention in the swine industry.

## Materials and methods

### Data

In this study, we used daily records of movement data of pigs in Upper Austria in 2021 and methods from social network analysis [22]. The Verbrauchergesundheitsinformationssystem (VIS) [26] is a national database where information on livestock movements and farms in Austria are recorded. Each pig movement is characterized by the date of the movement (dd.mm.yyyy), number of animals moved (batch size), type of movement (e.g., *domestic, slaughter, abroad*, and *abroad-slaughter*), source holding, and destination holding (more details on the data available in [25] and [27]). Each holding is identified via an anonymized unique identifier (ID). For each holding data includes information on the number of animals per production stage, the randomized 5 km-radius geocoordinates (i.e., latitude and longitude, projected coordinate system EPSG 31287), and the federal state where the holding was located (Upper Austria). Additionally, each holding is labeled according to its type of activity, as reported by the owner once a year. Labels include eight categories: abattoir, boar station, collection point, cutting plant, farm, private owner, processing plant, and trade and logistics. For this study, we excluded boar stations (i.e. production of quality liquid boar semen for artificial insemination), cutting and processing plants from the analysis since they were not involved in live animal movements and thus had limited importance in transmitting infectious diseases among live pigs. Holdings could have multiple labels if they reported different activities or no label when they did not report any activity for the year (the latter would hereafter be referred to as ”inactive” during the considered period).

The holdings’ geographical coordinates were overlaid over a shapefile of the Austrian municipalities [28] using the R package sf [29]. We extracted the name of the municipality where each holding was located using the function *st_ join()*. Due to the 5-kilometer radius randomization, some holdings fell outside the boundaries of Upper Austria. To address this, we employed a k-Nearest Neighbor join method (function *st_nn()*, with k = 1), which assigns the values of the closest polygon to a point, available in the package nngeo [30].

### Network analysis

To create the weighted networks, we assigned two types of edge weights: (i) trade frequency: each edge represents one trade with the weight 1, regardless of the number of pigs involved in the exchange; (ii) trade volume: each edge, representing one trade, is assigned a weight equal to the number of pigs traded during this exchange (i.e., batch volume). The edge direction corresponded to a movement *from* the source holding *to* the destination holding. We created two static representations of each network by aggregating the daily snapshots throughout 2021. We thereby obtained multigraphs, where multiple edges could exist between pairs of nodes, which were subsequently converted into simple graphs by collapsing the multiple edges into single-weighted ones, with edge weight corresponding to either trading frequency (i) or volume of animals traded (ii) between pairs of nodes. In both networks, a greater intensity in the weight of an edge between two nodes corresponded to a shorter “distance” between them, signifying a stronger connection. These two representations of the Upper Austrian pig trade network are isomorphic, meaning they share identical underlying structures and connections [31].

### Quantifying similarities and differences between networks

We normalized the edge weights in each network by dividing them by the maximum value of the edge weights within that specific network [32]. This normalization process was implemented to ensure that both networks were on a comparable scale, facilitating a fair comparison between them.

To characterize and compare both networks (frequency-versus volume-based), we computed the following weighted network centrality metrics, as suggested by Bellingeri *et al*. [22]: clustering coefficient [33], average path length [34], and diameter [34]. The clustering coefficient quantifies the density of triangles in a network, meaning the probability that two neighbors of the same node are themselves neighbors. The average path length is the average number of steps along the shortest paths for all possible pairs of nodes. The diameter is the length of the longest shortest path, i.e., among all shortest paths between every pair of nodes in the network for which a path exists [31, 33, 34].

Further, we calculated the following node centrality metrics: strength [7], betweenness [35], and closeness centrality [36]. Strength, which is the sum of the weights of the edges connected to a node, reflects the overall connectivity or influence of the node within the network. Betweenness centrality measures the extent to which a node lies on the shortest paths between other nodes, indicating its role as a bridge or connector within the network. While, closeness centrality is measuring the mean distance from a node to other nodes [1, 31, 35].

Closeness and betweenness centrality use shortest paths; however, in weighted networks of livestock moments, edge weights reflect the strength of the connection between a pair of holdings (nodes), not its cost. Therefore, to calculate weighted closeness and betweenness centrality, we inverted the edge weights, as proposed by Newman [37] and Brandes [38], ensuring that higher frequencies or volumes resulted in shorter distances.

The distributions of the three node centrality metrics and the edge weights in both networks were compared using density plots. To test if edges with high (low) weights were similar in both the frequency-based and the volume-based network, we ranked the edges based on their weight in each network and performed a Kendall rank correlation test with a significance level set to 0.05 [39]. Similarly, we performed a pairwise comparison of holding (node) rankings based on the three centrality metrics within the same network and between the frequency- and volume-based networks using the Kendall rank correlation test [39]. We computed the median centrality metrics for each type of activity and tested differences between groups using the Kruskal-Wallis test. Since a holding might have multiple activities, we included each multi-activity node in the calculations for each relevant label.

### Community detection

A community is defined as a group of nodes that have more connections (trades) to each other than with the rest of the network [40]. We used the Leiden algorithm, which encompassed three distinct steps: the initial step involved optimizing modularity, followed by the refinement of the partition, and finally, the third step focused on the community aggregation process [41]. First, to characterize trade communities among holdings we applied the Leiden algorithm on the above-described network of pig trades (holding level-community detection). Second, we built the network of pig trades at the municipality level by aggregating the holdings based on their municipality of location. This approach resulted in each node representing a municipality, with edges between nodes indicating pig movements between municipalities. Edge weights were assigned based on either the frequency of trades or the volume of pigs traded. The community detection algorithm was then run on the network of pig movements between Upper Austrian municipalities (municipality level).

We quantified the similarities in community membership using the matching method [42]. The matching value ranges from 0 to 1; a value of 1 signifies that the members of a community are identical, whereas if it is equal to 0, both communities display completely different members. We further compared the community patterns and the variation of community assemblages between both frequency-based and volume-based networks, in terms of holding/municipality composition, size, and spatial structure, including spatial connectivity.

### Epidemics on weighted networks

#### Spreading dynamic

We aimed to identify ”influential” nodes in the network, ranking them based on their ”reachability” during the spread of an epidemic. We define ”reachability” as the likelihood of a node to be infected during a contagion process. Specifically, we focused on identifying nodes that were infected more frequently than average during a contagion process. Additionally, we assessed whether node ranking, based on the contagion process, differs between frequency-based and volume-based networks and whether these results align with the rankings based on node centrality measures previously computed for both networks.

We simulated an epidemic spread on both networks using a Susceptible-Infectious-Recovered-Susceptible (SIRS) model. In this model, pig movements, represented by edges, indicate direct contact between pig operations, through which herds can be infected; each holding is assumed to belong to one of three distinct states: *S, I*, or *R*. A contact (trade) between an infectious source holding and a susceptible receiving one may lead to infection of the receiver. Infected holdings eventually recover after a certain period, during which they are considered empty holdings before restocking and resuming trade activities, thereby becoming susceptible again. The transition rate from I to R is denoted *γ*, and the transition rate from R to S is denoted *σ*. While transitioning from S to I, we introduced the edge weight, with edge types denoted as *ξ ∈ {*trade frequency, batch size*}*. As time progresses (*t* = 2, 3, …, *t*_*n*_), susceptible nodes *i* have a probability *λ*_*i*_ of being infected, with a transmission rate of *β* that depends on the weight of the edge *wξ* connecting them to infectious nodes. The probability *λ*_*i*_ is determined by summing the weights of the edges from all infectious nodes to node *i*.

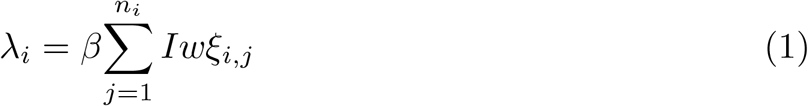

In this model, we set the values *β* = 0.5, *γ* ranged from 5 to 10 days, and *σ* ranged from 10 to 30 days. We selected the *β* value based on Büttner *et al*. [3], where high transmission probability led to a convergence in the number of infected farms across differently weighted networks. The range of *γ* was chosen to allow sufficient time for the infection to spread. Finally, *σ* was determined based on the scientific opinion of the European Food Safety Authority (EFSA) Panel on Animal Health and Welfare [43].

Although the epidemic spread was simulated at the holding level, the initial infected holding was chosen randomly from a randomly selected municipality within each previously identified community. This approach, based on the results of the community detection performed at holding versus municipality level (see S1 Text), ensured that the seeding of the infection occurs with equal probability across all communities. Nodes labeled exclusively as abattoirs were excluded from the seeding process. At the onset of the epidemic (*t* = 1), an initially infected holding was introduced. For each network representation and every community (where the epidemic could start), 1000 simulations were run iteratively for *t* = 90 days.

We evaluated the overall impact and breadth of the epidemic by calculating the average number of infected holdings over all simulations (epidemic size) and the average number of communities reached by the epidemic (S2 Text).

#### Node ranking based on nodes’ reachability

We counted the number of times each node *i* was infected or reinfected within a single simulation. For each node *i* in community *C*_*j*_, we constructed a vector *V*_*i,j*_ = (*n*_*j*1_, …, *n*_*j*1000_), where each element *n*_*jk*_ represents the number of times node *i* was (re)infected in the *k*-th simulation, with *i ∈ {*1, number of nodes*}, j ∈ {*1, number of communities*}*, and *k ∈ {*1, 1000*}*. Next, we computed the average infection frequency for each node in both networks’ representation across all simulations. This average, denoted as *Ii*, represents the number of times the epidemic passes through *i* across all simulations. Using *Ii*, we ranked the nodes based on their reachability during a contagion process, which we termed as simulation-based ranking. This simulation-based ranking served as the benchmark for comparison with centrality-based rankings.

After obtaining the simulation-based ranking, we compared it with centrality-based rankings using Kendall rank correlation tests. Nodes in the top 5% according to the highest correlated centrality metric were considered potential candidates for sentinel surveillance holdings. We analyzed their spatial distribution using point pattern analysis [44] to assess clustering or random spatial distribution. For further details on the methods see S3 Text.

#### Software

Data cleaning, preparation, subsequent data analyses, and computation of the figures were performed in R software (v.4.1.0) [45] using RStudio Server ”Juliet Rose” (v.1.4.1717) [46]. The network analysis and study of the spreading dynamics was performed using the R package igraph [47]. Community detection using the Leiden algorithm was conducted using leidenAlg [48]. Point pattern analysis was performed with spatstat [49].

## Results

### The pig trade network in Upper Austria

In 2021, there were 110,891 movements recorded in Upper Austria involving a total of 6,020 pig holdings. Of these, 254 (4.2%) engaged in animal trade with other Austrian federal states or countries. For this study, we selected movements that occurred only within the federal state of Upper Austria, accounting for 92,914 movements (83.8%) involving 5,766 holdings (95.8%). This included 5,726 (99.3%) farms, 288 (5%) abattoirs, 96 (1.6%) trading points, and 14 (¡1%) collection points. Overall, the data represents a volume of 3,100,852 pigs traded, of which, 1,510,690 (49%) were intended for slaughter while 1,590,162 (51%) were transferred between animal operations.

The majority of the holdings were concentrated in central Upper Austria, specifically in the municipalities of Vorchdorf and Pettenbach as well as in the northeastern region, including the municipality of KÖnigswiesen (Fig 1).

**Fig 1.**
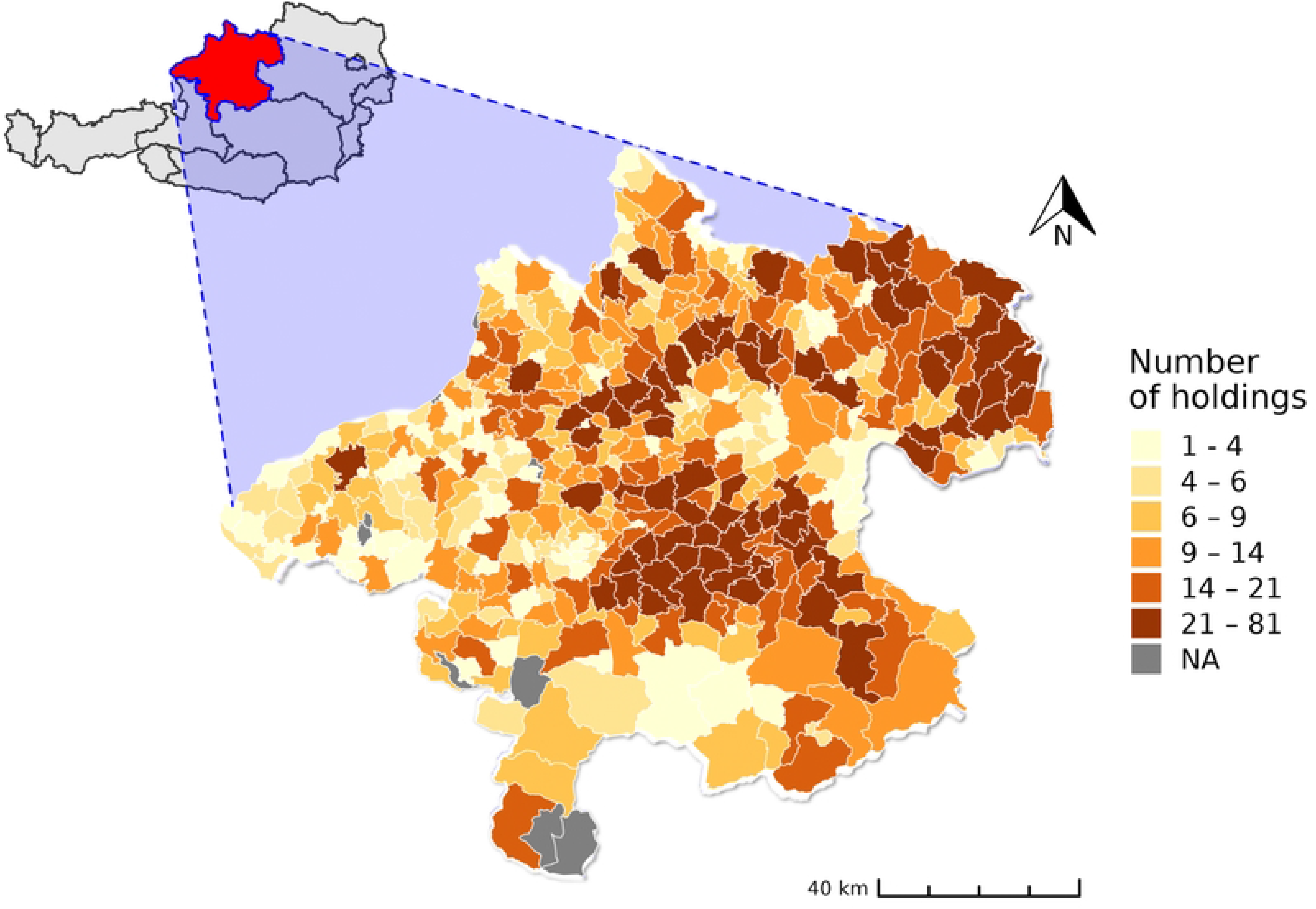
Density of pig holdings per municipality in the federal state of Upper Austria, Austria, 2021. KÖnigswiesen (in the northeast), Vorchdorf, and Pettenbach (in the center) exhibit the highest density of holdings.

### Comparison of the edge weight distributions

In the frequency-based pig trade network, the trade frequency between pairs of pig operations varied from 1 to 99, with a median of 2. In the volume-based network, the edge weights, corresponding to the total number of pigs exchanged between two holdings, ranged from 1 to 9,384 animals, with a median of 26. In this network, 896 edges (5.5%) accounted for half of the total volume of animals transferred. Overall, these figures revealed that the majority of pig holdings had infrequent and low-volume exchanges during the studied period. In contrast, a small number of trade partners exhibited sustained connections with high transaction frequency and volume.

The edge weight distribution patterns differed between the frequency-based and volume-based networks. Specifically, the edge weight distribution in the frequency-based network showed a clear bimodal pattern, whereas the edge weight distribution in the volume-based network exhibited a more uniform pattern (Fig 2). The Kendall test showed a strong positive correlation in the ranking of edges based on their weights between both networks (Kendall tau-b (*τb*) correlation coefficient = 0.56; *p <* 0.001), suggesting consistent ordering of edge weights across both networks.

**Fig 2.**
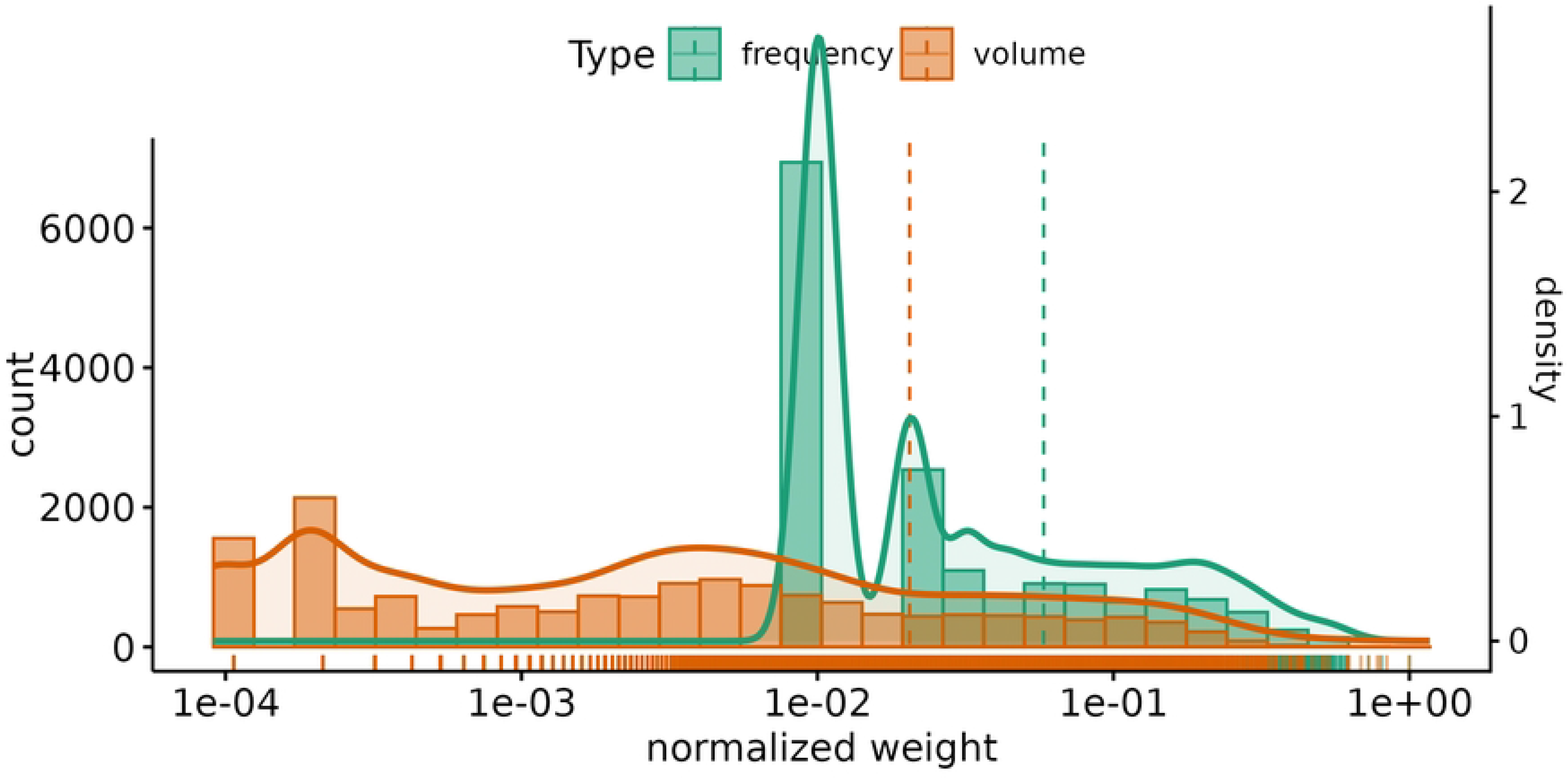
Edge weight distribution in the frequency-(green) and volume-based network (red) of pig trade in Upper Austria, 2021, normalized, in each network, by the maximum edge value. The dashed lines and marginal rugs represent the means and marginal distribution for each network, respectively. The x-axis is presented in a log scale.

### Network centrality statistics

The clustering coefficients of both networks were similar, 8.9 *×* 10^*−*3^ for the frequency-based network versus 8.7 *×* 10^*−*3^ for the volume-based network. The average path length of the frequency-based network was 1.6, while it was 0.3 in the volume-based network, indicating that holdings were connected by less than one or two links (trades) on average, respectively. The diameter of the frequency-based network was 6.97, whereas it was smaller (4.09) in the volume-based network. These findings suggested that the frequency-based network was more densely connected than the volume-based network.

**Table 1.** Network-level measures calculated for the frequency- and volume-based representations of the Upper Austrian pig trade network, Austria, 2021.

| Metric | Frequency-based | Volume-based |
| --- | --- | --- |
| Clustering coefficient | $8.9 \times 10^{-3}$ | $8.7 \times 10^{-3}$ |
| Average path length | 1.6 | 0.3 |
| Diameter | 6.97 | 4.09 |

Clustering coefficient, average path length, and diameter for both the frequency- and volume-based pig trade networks in Upper Austria, Austria, 2021.

### Community detection

#### Holding-level community detection

The Leiden algorithm identified 46 and 80 trade communities within the frequency-based and volume-based networks of pig trades between Upper Austrian holdings, respectively. These communities were characterized by high spatial fragmentation, often comprising fewer than five holdings each (see S1 Text). This output was not further considered in this study, as the focus was on larger, more interconnected communities that could offer meaningful insights into pig trade dynamics and network structures, particularly for epidemiological purposes.

#### Municipality-aggregated network

The Leiden algorithm identified six trade communities within the frequency-based and volume-based networks of pig trades between Upper Austrian municipalities (Fig 3).

**Fig 3.**
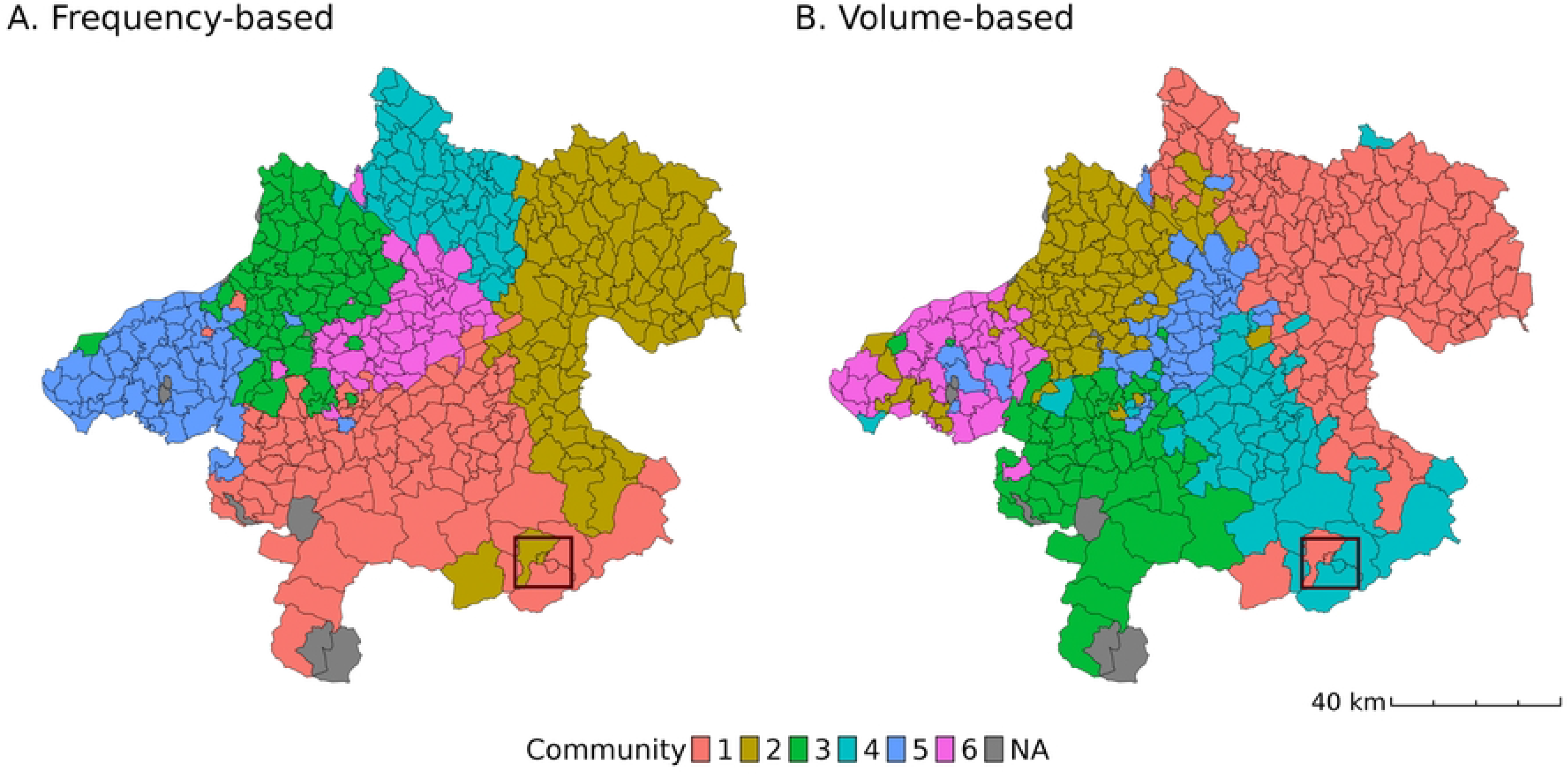
Pig trade communities in Upper Austria, Austria, 2021, detected using the Leiden algorithm run on the municipality-aggregated network. A) Frequency-based network. B) Volume-based network. The municipality is considered as an epidemiological unit. The black square shows where the municipality of Windischgarsten is located.

The matching coefficient was 0.22*±*0.29 (standard deviation, SD) indicating that the community memberships in both network representations differed. The frequency-based network displayed distinct community boundaries (Fig 3A), indicating spatial segregation among different communities. Conversely, the volume-based network exhibited more fragmented communities, i.e., the communities did not form distinct ”blocks” (Fig 3B). Specifically, while certain regions consistently showed membership in the same communities across both networks, particularly in the eastern and southern parts of Upper Austria, other areas in the north, center, and west exhibited varying community memberships. Spatially isolated municipalities belonging to a distanced community typically had only one edge connecting them to a member of that community. For example, the municipality Windischgarsten in the south of Upper Austria (black square in Fig 3) was connected to the municipality Rossleithen, also in the south, by a single trade involving ten animals.

### Comparison of node rankings

#### Ranking based on node centrality metrics

We first compared node rankings based on weighted centrality metrics within each network (Fig 4). The Kendall’s *tau* revealed strong correlations between node rankings based on strength and closeness for both networks (*τb*= 0.75 for the frequency-based network and *τ* = 0.84 for the volume-based network; *p <* 0.001). The correlation between rankings derived from closeness and betweenness in both networks exhibited a lower but significant correlation *τb*= 0.35; *p <* 0.001 for both networks. Node rankings based on strength and betweenness displayed moderate correlations in both networks (*τb*= 0.47 for the frequency-based network and *τ* = 0.42 for the volume-based network; *p <* 0.001). These findings underscored that betweenness is weakly correlated with the two other measures in both networks.

**Fig 4.**
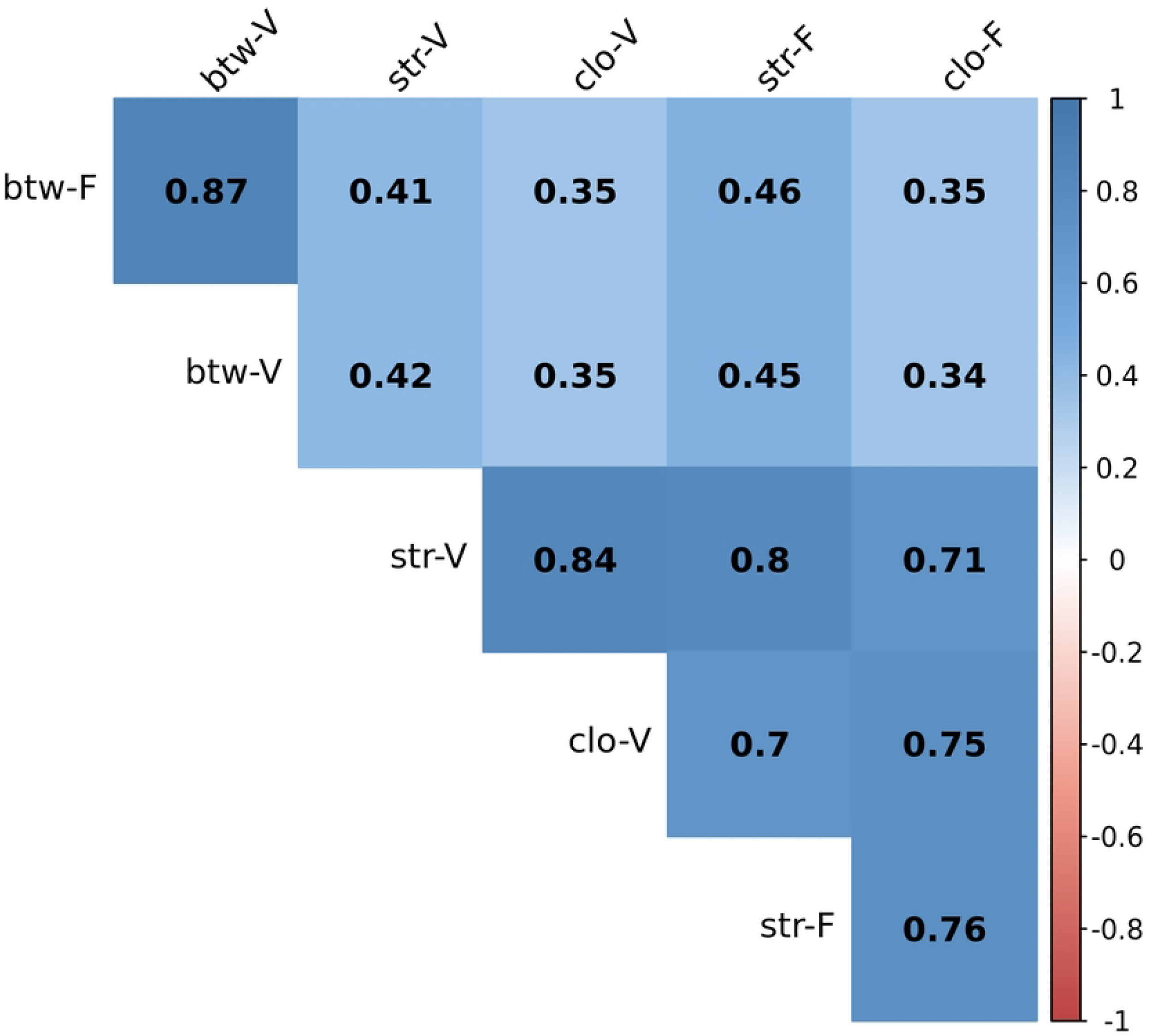
Upper matrix of pairwise Kendall’ s-tau rank correlations between node rankings based on three weighted centrality metrics (strength; str; closeness: clo; betweenness: btw) within and between the frequency-based (denoted F-) and volume-based (V-) representations of the Upper Austrian pig trade network, 2021. The Kendall-tau correlation coefficient can take values from -1 to 1, where the most positive value reflects a positive correlation between ranks.

When comparing the two networks, a strong correlation (*τb*= 0.8, *p <* 0.001) was found between rankings based on node strength. Among the top 5% nodes (representing 289 holdings) with the highest strength centrality in both frequency- and volume-based networks, 107 nodes were common in both. The density plots of the node strength distributions (Fig 5A) revealed a bimodal pattern. A notable proportion of nodes showed lower strength centrality values in the volume-based network, while most of the nodes in the frequency-based had average values.

**Fig 5.**
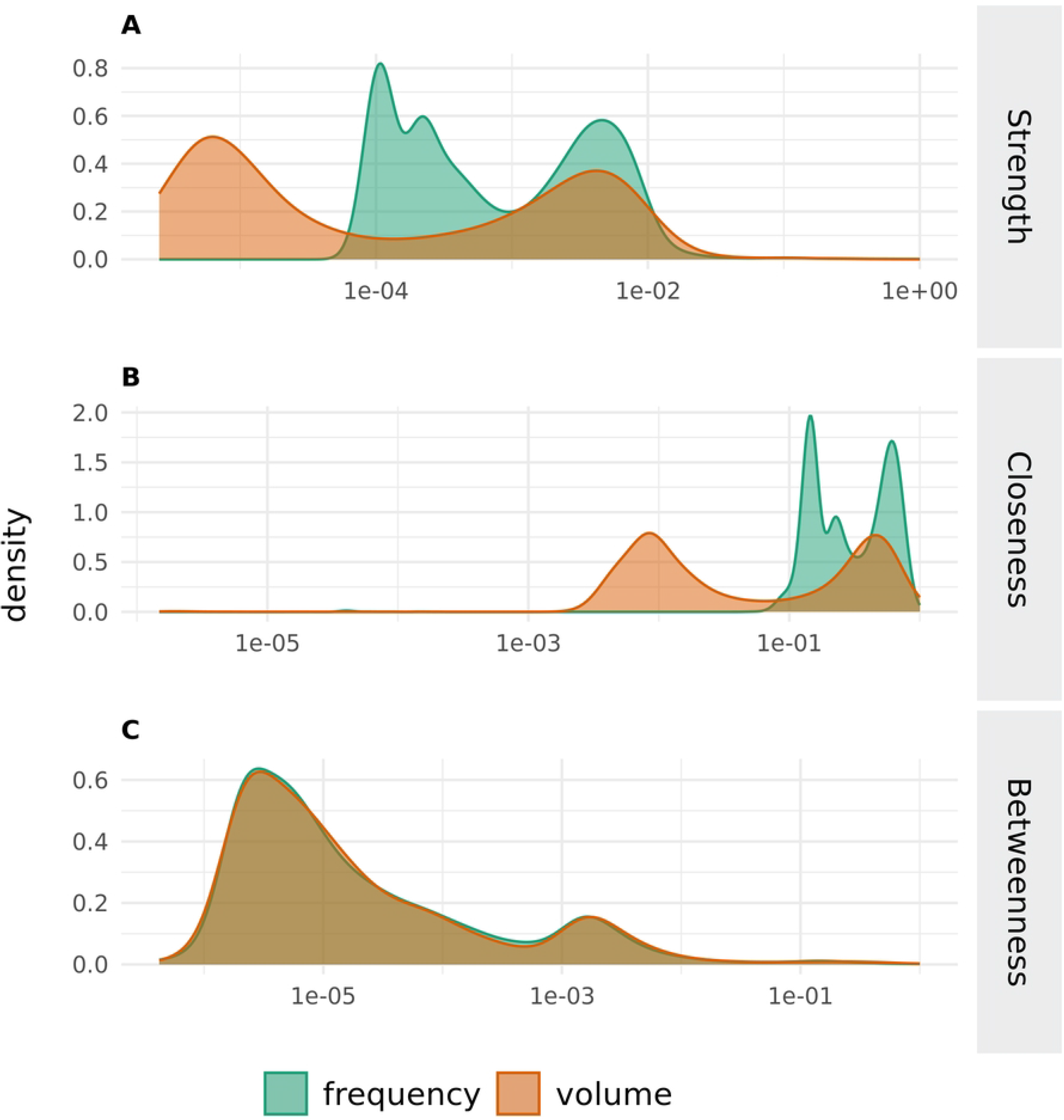
Density plots showing the distributions of the weighted node centrality metrics A) strength, B) closeness, and C) betweenness for both the frequency- and volume-based networks of pig movements in Upper Austria, Austria, 2021.

Similarly, correlation analysis of node rankings based on closeness centrality between the networks revealed a strong correlation (*τb*= 0.75, *p <* 0.001). We identified 148 common nodes to the top-ranked nodes in both networks. The density plots of the node closeness centrality distributions resembled those of the node strength distributions, exhibiting a bimodal pattern with some overlap (Fig 5B).

Additionally, correlation analysis of node rankings based on betweenness centrality between the networks also revealed a strong correlation (*τb*= 0.86, *p <* 0.001). Among the top-ranked nodes in both networks based on betweenness, 244 nodes overlapped. The density plot of the node betweenness distributions displayed a bimodal pattern, although skewed towards lower values, with a high degree of overlap, confirming a highly similar distribution of node betweenness in both networks (Fig 5C).

The Kruskal-Wallis test indicated statistically significant differences in centrality metrics between pig operation types (*p <* 0.001). In both networks, animal operations labeled as ”collection points” consistently showed high median centrality values across all three metrics. On the other hand, slaughterhouses demonstrated high median strength and closeness centrality but lower betweenness centrality values (Tab 2). The role of slaughterhouses as endpoints for live pigs contributes to their low betweenness centrality, meaning they do not typically lie on a path between other nodes. Farms exhibited generally lower median centrality metrics because a large number of farms demonstrated a relatively low trade frequency and lower volume of pig transferred compared to other pig operations (e.g., collection points frequently received and sent a large amount of pigs, abattoirs received frequently large volume of animals for slaughter).

**Table 2.**
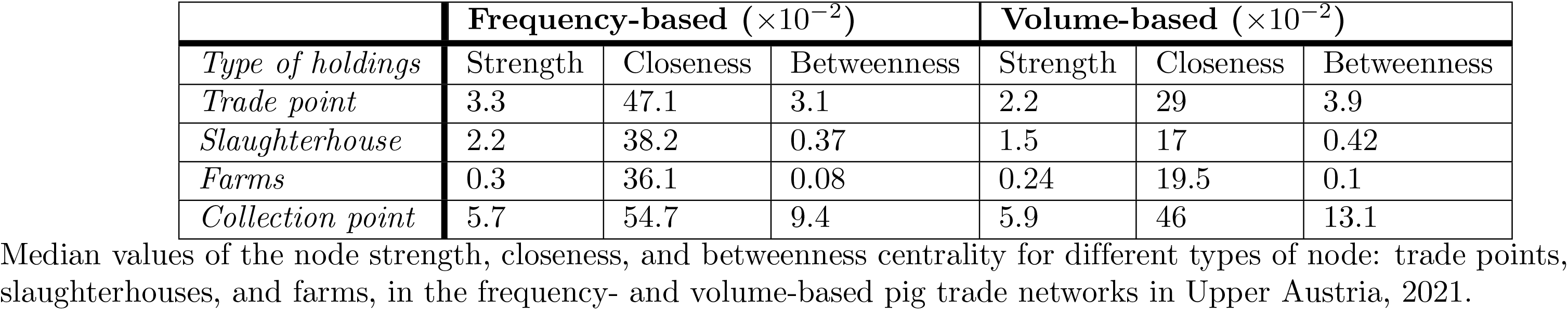
Node centrality values for both the frequency- and volume-based networks of pig movements in Upper Austria, Austria, 2021.

### Rankings of nodes based on nodes’ reachability during an epidemic

We seeded the infection an equal number of times in each community to ensure an unbiased probability of initial infection location and to assess whether the epidemic would remain within individual communities or spread throughout the entire network. We observed that the disease propagation process extended beyond the initially seeded community, reaching nodes in other communities within both the frequency-based and volume-based networks (S2 Text). In the subsequent analysis, we aggregated the results from 6000 simulations initiated across these six communities.

### Comparing the simulation-based rankings and epidemic size

In the frequency-based network, a node was infected, on average, 0.022 times per simulation (95%CI: 0.02-0.023); this measure of reachability was slightly lower in the volume-based network at 0.017 (95%CI: 0.016-0.018). We further assessed the agreement between network rankings using Kendall’s tau-b, and revealed a strong positive correlation (*τb*=0.678, *p <* 0.001) between the simulation-based rankings of the two networks. This indicates a high consistency in node importance assessed through simulation analysis of an epidemic propagation model across both network representations.

The epidemic size (i.e., number of nodes infected at least once, averaged over 1000 simulations) based on the municipality of initial infection is reported in S2 Text. Results of the non-parametric Kruskall test showed significant differences (*p <* 0.005) in the epidemic size depending on the community chosen for the initial seeding of the epidemic in both network representations. Post-hoc analysis with Dunn’s multiple comparisons test revealed that in the frequency-based network, seeding in Community 6 led to the greatest facilitation of epidemic spread (i.e., greatest epidemic size), while Community 5 showed the least facilitation. In the volume-based network, seeding in Community 4 consistently facilitated the highest level of epidemic spread, while Community 6 demonstrated the lowest facilitation. Additionally, the Wilcoxon test showed a significant difference in the median epidemic size between the frequency-based and volume-based networks (*p <* 0.005). Furthermore, communities 5 and 3 consistently exhibited the highest vulnerability to epidemics (as measured by the percentage of infected holdings per community), regardless of the seeding point, in both the frequency-based and volume-based network.

### Comparing centrality-based and simulation-based rankings

We compared the simulation-based rankings of nodes in both networks with rankings based on each centrality measure (i.e., strength, closeness, and betweenness). In both the frequency-based and volume-based networks, the simulation-based ranking showed weak positive correlations with rankings based on the three centrality measures 3. In both network representations, node ranking based on strength centrality exhibited the highest correlation with the simulation-based ranking. Further analysis of the top 5% of nodes (289 nodes) for each ranking revealed a stronger positive correlation between strength centrality-based and simulation-based rankings compared to the entire node list. In contrast, the *τb* values for rankings based on closeness centrality indicated lower agreement with the simulation-based ranking for the top 5%.

Since the rank based on strength centrality showed the strongest correlation in both network representations, we focused our subsequent analysis on the top 5% highest-ranked nodes based on this metric, and evaluated their potential use as sentinels in surveillance programs. In the frequency-based network, the top 5% highest-ranked nodes based on strength centrality included 113, 50, 45, 16, 8, and 57 nodes distributed across communities 1 to 6, respectively. Similarly, in the volume-based network, the top 5% highest-ranked nodes based on strength centrality comprised 107, 14, 84, 31, 15, and 38 nodes located in communities 1 to 6, respectively. The number of nodes common in the highest-ranked nodes for both networks was 107 nodes. We examined the spatial distribution of these potential sentinels in both networks using point pattern analysis. The results indicated that these nodes were spatially clustered in both networks as compared to what would be expected under Complete Spatial Randomness (Figure 3 in S3 Text) (*p − values <* 0.005). Furthermore, when comparing the spatial distribution of these potential sentinels between both networks, we evidenced that potential sentinel holdings in the volume-based network exhibited tighter clustering patterns compared to the potential sentinels holdings identified in the frequency-based network (S3 Text).

## Discussion

In this work, we employed various approaches to compare two isomorphic networks of animal movements weighted differently based on trade frequency or volume of animals traded. We examined network properties and community structures, and explored how edge weights influence node importance using three centrality metrics and a measure of reachability. While both networks shared similarities in terms of the number of detected communities and node rankings derived from centrality metrics and simulations, we demonstrated that the choice of edge weight significantly affects edge weight distributions, community memberships, and epidemic size.

**Table 3.** Comparison of node rankings based on three centrality metrics versus simulation-based rankings, for both the frequency- and volume-based networks of pig movements in Upper Austria, Austria, 2021.

|  |  | Full network |  | Top 5% highest-ranked |  |
| --- | --- | --- | --- | --- | --- |
| Network | Centrality | tau | p-value | tau | p-value |
| Frequency-based | betweenness | 0.153 | < 0.001 | 0.076 | 0.3387 |
|  | closeness | 0.133 | < 0.001 | 0.08 | 0.3436 |
|  | strength | 0.155 | < 0.001 | 0.51 | < 0.001 |
| Volume-based | betweenness | 0.151 | < 0.001 | 0.2 | 0.02396 |
|  | closeness | 0.167 | < 0.001 | -0.024 | 0.7753 |
|  | strength | 0.173 | < 0.001 | 0.5 | < 0.001 |

Kendall’s tau coefficients and associated p-values from comparing node rankings based on betweenness, closeness, and strength centrality with node reachability determined through simulated epidemic spread in frequency-based and volume-based networks.

While the edge weight distribution in the frequency-based network exhibited a clear bimodal pattern, in contrast to a more uniform distribution in the volume-based network, we found a strong correlation in the ranking of edges based on their weight. This suggests that the underlying trade patterns were similar across both network representations, i.e., pairs of holdings engaging in frequent exchanges also traded higher volumes of animals. Therefore, focusing on the top-ranked edges based on their weights, regardless of the network representation, is crucial for identifying critical connections relevant to disease control.

Our findings revealed a strong positive correlation among node rankings based on centrality metrics (strength, closeness, and betweenness) both between the two network representations and within each network. This ranking agreement suggested that multiple centrality measures may redundantly capture similar aspects of node importance within a single network [50]. Moreover, high correlations across networks with different weights indicated that both networks shared similar underlying properties.

However, betweenness centrality consistently exhibited lower correlations with closeness and strength centrality in both networks. This discrepancy may arise because betweenness centrality identifies nodes that serve as bridges connecting different parts of the network, whereas closeness and strength centrality emphasize nodes with central locations and strong connections, respectively [31, 51].

Our results highlight that collection points consistently showed high node centrality values across all three metrics in both network representations, indicating their central role in the networks, irrespective of edge weight, which underscored a core property of the studied pig trade network. These holdings facilitate numerous transactions, functioning as bridges [51]. On the other hand, farms and slaughterhouses in both networks exhibited low betweenness centrality values, emphasizing their roles at the end of pork production. Indeed, pigs typically originate from many farms and then move unidirectionally and convergently toward a fewer number of slaughterhouses.

Ranking nodes based on their average reachability (the average number of times a node is reached by an epidemic over 6000 simulations) revealed a strong correlation between the two networks. This suggested that edge weights have minimal impact on node reachability. In other words, the identification of vulnerable nodes during an epidemic appears consistent regardless of whether the analysis considered the frequency or volume of interactions between nodes.

Strength centrality, which considered both the number of connections (degree centrality) and their weights (intensities) [31, 51], exhibited the strongest correlation with the simulation-based ranking of influential nodes in both networks, i.e., independently of the weight used. This correlation becomes stronger when focusing on the top 5% highest-ranked nodes. This makes strength centrality a valuable tool for identifying vulnerable holdings in the network, i.e., those frequently involved in disease transmission. Our findings align with those of Candeloro et.al [52], who demonstrated that their newly defined weighted strength and degree centrality measures (accounting for edge weights distribution) best capture nodes importance in the spread of epidemic disease and can be utilized to identify influential nodes in the network that can be used for targeted surveillance programs.

The weaker correlation observed across the full network is due to the instability of the rankings’ tails, influenced by a large number of holdings that infrequently engaged in trades or involved relatively small volumes of animals. Similar observations of pig operations with sporadic trading patterns have been documented in other pig trade networks [3, 8–10].

Our research identified a significant structural divergence in network behavior based on data aggregation level, evident in both frequency-based and volume-based network representations. Specifically, at the holding level, the network demonstrated low modularity, whereas the municipality-aggregated network exhibited high modularity. Notably, our analysis of the top 5% influential holdings by strength centrality revealed a clustered spatial distribution across Upper Austria. However, this distribution was uneven across communities within the municipality-aggregated network, which further supports the use of the top 5% highest-ranked nodes as effective sentinels in surveillance programs. This finding aligns with insights from Colman et.al [53], who proposed that for highly modular networks, such as observed for the municipality-aggregated network, sentinel surveillance strategies are most effective when sampling efforts are dispersed across different regions or communities.

Our results emphasize significant differences in the epidemic size depending on the chosen community for the initial seeding of the epidemic in both network representations. Moreover, the average epidemic size was significantly higher in the frequency-based network compared to the volume-based network. Epidemic spread may be facilitated in some communities due to the presence of pairs of holdings with weak ties that act as bridges between communities, exemplifying the ’strength of weak ties’ paradox [54, 55]. weak ties, characterized by infrequent or low-volume exchanges, serve as connectivity-generating factors and tend to function as crucial connectors that link otherwise distant communities. As a result, they extend connections beyond their own community, facilitating the spread of phenomena, including diseases.

This study presents some limitations. First, it focuses on the pig trade network in Upper Austria, thus the results may only be representative of this specific regional trade network. Additionally, we limited our analysis to three centrality metrics. While these metrics are widely used in similar network analyses [9, 22, 23, 53, 56–58], exploring other centrality metrics might reveal further consensus with the simulation-based ranking, particularly for networks with characteristics similar to those of the Upper Austria pig trade network, specifically those exhibiting the small-world properties [59].

Second, we used a single transmission rate for the disease spreading simulations. Conducting additional simulations with varying transmission rates, particularly lower ones, could yield different results for node reachability rankings and enhance our understanding of how different transmission dynamics affect these rankings. For example, Büttner et al. [3] found that transmission probabilities below 0.5 led to significant differences in the number of infected farms between networks weighted by trade contacts or number of delivered pigs. Notably, they demonstrated that, as transmission probability increases, the frequency-weighted network experiences a higher number of infected farms. Additionally, incorporating additional information about holdings, such as biosecurity levels and operational sizes, could further refine the model and may affect the node ranking based on reachability.

Moreover, this study used a static representation of the pig trade network. Static network models, while providing an overview of network structure, may not accurately reflect the dynamic nature of network topology over time. This meant our static centrality measures and epidemic simulations on aggregated weighted networks might not have fully captured temporal variations in trades, potentially overlooking fluctuations or seasonal changes throughout the year [9]. From an epidemiological standpoint, this implies that while some infectious contacts between holdings may be long-term, many are transient [60].

However, the practicality of a surveillance strategy based on node strength centrality must be considered. If rankings based on centrality metrics change over time, stakeholders may face challenges in adapting their surveillance efforts to these evolving rankings. Nevertheless, increasing network analysis complexity could hinder field workers’ ability to effectively implement findings. Therefore, while dynamic network representations are necessary for future research [61], the usability of these models in the field should also be evaluated to ensure they remain manageable for those performing surveillance.

## Conclusion

Our findings have practical implications for developing surveillance strategies. The strong alignment of ranking derived from strength centrality with simulation-based ranking suggests that this metric can be used to identify critical nodes in the network, potentially avoiding the need for computationally demanding and time-consuming simulations. Network-based approach can enhance the efficiency of traditional surveillance strategies, making them more effective and field-deployable.

Both network representations exhibited considerable structural similarity, although the choice of the edge weighting method significantly influences epidemic size. This underscores the importance of tailoring edge weight selection to the specific characteristics of the infectious disease under surveillance or being modeled, notably its transmission probability. Such a tailored approach ensures that surveillance efforts are optimally aligned with the disease transmission dynamics within the network.

In conclusion, the weighted network approach shows potential for developing targeted surveillance strategies. By focusing on critical nodes identified through centrality measures, particularly strength centrality, and understanding the differences between frequency-based and volume-based network representations, we can develop more efficient and cost-effective disease control measures.

## Supporting information

**S1 Text.Holding-level community detection**. Results from the Leiden algorithm applied to the network of pig trades in Upper Austria, 2021, detailing the number of communities detected and the member count within each community.

**S2 Text. Disease spread within and across communities**. Analysis of epidemic spread dynamics to determine whether outbreaks remain within the initial community or extend to other communities.

**S3 Text.Point pattern analysis**. Analysis of the spatial distribution of the top 5% highest-ranked nodes based on strength centrality.

## Data and code availability

The metadata and R code used to produce the results of this study will be made publicly available on Figshare upon publication. The raw data that support the findings of this study are available from the Austrian Federal Ministry of Social Affairs, Health, Care and Consumer Protection (BMSGPK) but restrictions apply to the availability of these data, which were used under license for the current study, and so are not publicly available. Data are however available from the authors upon reasonable request and with permission of the data owner.

## Acknowledgments

The authors would like to thank the Austrian Federal Ministry of Social Affairs, Health, Care and Consumer Protection (BMSGPK) and the Austrian Agency for Health and Food Safety (AGES) for providing the data for this study. We also extend our gratitude to Andrea Ladinig and Klemens Fuchs, members of our advisory board, for their expert feedback and professional guidance. We also want to thank the Infectious Disease and One Health Erasmus Mundus Joint Master Degree for facilitating the internship of Yan-Shin Jackson Liao in our Group. Finally, we are grateful to our colleagues at the Complexity Science Hub for their valuable brainstorming sessions and continuous support.

## Notes

### Competing Interest Statement

The authors have declared no competing interest.

## References

1. Keeling MJ, Rohani P. Modeling Infectious Diseases in Humans and Animals. Princeton University Press; 2011.

2. Newman MEJ. Analysis of weighted networks. Physical Review E. 2004;70:056131. doi:10.1103/PhysRevE.70.056131.

3. Bºttner K, Krieter J. Epidemic spreading in a weighted pig trade network. Preventive Veterinary Medicine. 2021;188:105280. doi:10.1016/j.prevetmed.2021.105280.

4. Jolly AM. Sexual Networks and Sexually Transmitted Infections: A Tale of Two Cities. Journal of Urban Health: Bulletin of the New York Academy of Medicine. 2001;78:433–445. doi:10.1093/jurban/78.3.433.

5. Lockhart AB, Thrall PH, Antonovics J. Sexually Transmitted Diseases In Animals: Ecological and Evolutionary Implications. Biological Reviews. 1996;71:415–471. doi:10.1111/j.1469-185X.1996.tb01281.x.

6. Kamp C, Moslonka-Lefebvre M, Alizon S. Epidemic Spread on Weighted Networks. PLoS Computational Biology. 2013;9:e1003352. doi:10.1371/journal.pcbi.1003352.

7. Barrat A, Barth©lemy M, Pastor-Satorras R, Vespignani A. The architecture of complex weighted networks. Proceedings of the National Academy of Sciences. 2004;101:3747–3752. doi:10.1073/pnas.0400087101.

8. Passafaro TL, Fernandes AFA, Valente BD, Williams NH, Rosa GJM. Network analysis of swine movements in a multi-site pig production system in Iowa, USA. Preventive Veterinary Medicine. 2020;174:104856. doi:10.1016/J.PREVETMED.2019.104856.

9. Lentz HHK, Koher A, Hvel P, Gethmann J, Sauter-Louis C, Selhorst T, et al. Disease Spread through Animal Movements: A Static and Temporal Network Analysis of Pig Trade in Germany. PLOS ONE. 2016;11:e0155196. doi:10.1371/JOURNAL.PONE.0155196.

10. O’Hara KC, Beltr°n-Alcrudo D, Hovari M, Tabakovski B, Martnez-Lpez B. Network analysis of live pig movements in North Macedonia: Pathways for disease spread. Frontiers in Veterinary Science. 2022;9. doi:10.3389/fvets.2022.922412.

11. Schley D, Whittle S, Taylor M, Kiss IZ. Models to capture the potential for disease transmission in domestic sheep flocks. Preventive Veterinary Medicine. 2012;106:174–184. doi:10.1016/j.prevetmed.2012.01.023.

12. Begon M, Bennett M, Bowers R¬, French N¬, Hazel S¬, Turner J. A clarification of transmission terms in host-microparasite models: numbers, densities and areas. Epidemiology and Infection. 2002;129:147–153. doi:10.1017/S0950268802007148.

13. McCallum H. How should pathogen transmission be modelled? Trends in Ecology Evolution. 2001;16:295–300. doi:10.1016/S0169-5347(01)02144-9.

14. Anderson RM, May RM. Population biology of infectious diseases: Part I. Nature. 1979;280:361–367. doi:10.1038/280361a0.

15. Bouma A, de Jong MCM, Kimman TG. Transmission of pseudorabies virus within pig populations is independent of the size of the population. Preventive Veterinary Medicine. 1995;23:163–172. doi:10.1016/0167-5877(94)00442-L.

16. Chin WCB, Bouffanais R. Spatial super-spreaders and super-susceptibles in human movement networks. Scientific Reports. 2020;10(1):18642. doi:10.1038/s41598-020-75697-z.

17. da Silva RAP, Viana MP, da Fontoura Costa L. Predicting epidemic outbreak from individual features of the spreaders. Journal of Statistical Mechanics: Theory and Experiment. 2012;2012(07):P07005. doi:10.1088/1742-5468/2012/07/P07005.

18. Kitsak M, Gallos LK, Havlin S, Liljeros F, Muchnik L, Stanley HE, et al. Identification of influential spreaders in complex networks. Nature Physics. 2010;6(11):888–893. doi:10.1038/nphys1746.

19. Albert R, Jeong H, Barab°si AL. Diameter of the World-Wide Web. Nature. 1999;401(6749):130–131. doi:10.1038/43601.

20. Bºttner K, Krieter J, Traulsen A, Traulsen I. Static network analysis of a pork supply chain in Northern GermanyÄîCharacterisation of the potential spread of infectious diseases via animal movements. Preventive Veterinary Medicine. 2013;110:418–428. doi:10.1016/j.prevetmed.2013.01.008.

21. Rautureau S, Dufour B, Durand B. Structural vulnerability of the French swine industry trade network to the spread of infectious diseases. Animal. 2012;6:1152–1162. doi:10.1017/S1751731111002631.

22. Considering weights in real social networks: A review. Frontiers in Physics. 2023;11. doi:10.3389/fphy.2023.1152243.

23. Bucur D, Holme P. Beyond ranking nodes: Predicting epidemic outbreak sizes by network centralities. PLOS Computational Biology. 2020;16:e1008052. doi:10.1371/journal.pcbi.1008052.

24. Statistics Austria. Austria. Figures. Data. Facts; 2024. Available from: www.statistik.at.

25. Puspitarani GA, Fuchs R, Fuchs K, Ladinig A, Desvars-Larrive A. Network analysis of pig movement data as an epidemiological tool: an Austrian case study. Scientific Reports. 2023;13(1):9623. doi:10.1038/s41598-023-36596-1.

26. Statistics Austria. Statistics Austria;.

27. Puspitarani GA, Fuchs R, Fuchs K, Ladinig A, Desvars-Larrive A. Network analysis of pig movement data as an epidemiological tool: an Austrian case study; 2023. Available from: 10.6084/m9.figshare.21904995.v1.

28. Statistik Austria. Gliederung nsterreichs in Gemeinden -Dataset - data.gv.at;.

29. Pebesma E. Simple Features for R: Standardized Support for Spatial Vector Data. The R Journal. 2018;10(1):439–446. doi:10.32614/RJ-2018-009.

30. Dorman M. nngeo: k-Nearest Neighbor Join for Spatial Data; 2023. Available from: https://CRAN.R-project.org/package=nngeo.

31. Wasserman S, Faust K. Social Network Analysis. Cambridge University Press; 1994. Available from: https://www.cambridge.org/core/product/identifier/9780511815478/type/book.

32. Antoniou IE, Tsompa ET. Statistical Analysis of Weighted Networks. Discrete Dynamics in Nature and Society. 2008;2008:1–16. doi:10.1155/2008/375452.

33. Mcassey MP, Bijma F. A clustering coefficient for complete weighted networks. Network Science. 2015;3:183–195. doi:10.1017/nws.2014.26.

34. West DB, et al. Introduction to graph theory. vol. 2. Prentice hall Upper Saddle River; 2001.

35. Freeman LC. Centrality in social networks conceptual clarification. Social Networks. 1978;1:215–239. doi:10.1016/0378-8733(78)90021-7.

36. Marchiori M, Latora V. Harmony in the small-world. Physica A: Statistical Mechanics and its Applications. 2000;285:539–546. doi:10.1016/S0378-4371(00)00311-3.

37. Newman MEJ. Scientific collaboration networks. II. Shortest paths, weighted networks, and centrality. Physical Review E. 2001;64(1):016132. doi:10.1103/PhysRevE.64.016132.

38. Brandes U. A faster algorithm for betweenness centrality*. The Journal of Mathematical Sociology. 2001;25(2):163–177. doi:10.1080/0022250X.2001.9990249.

39. Ghanem M, Magnien C, Tarissan F. Centrality Metrics in Dynamic Networks: A Comparison Study. IEEE Transactions on Network Science and Engineering. 2019;6:940–951. doi:10.1109/TNSE.2018.2880344.

40. Radicchi F, Castellano C, Cecconi F, Loreto V, Parisi D. Defining and identifying communities in networks. Proceedings of the National Academy of Sciences. 2004;101(9):2658–2663. doi:doi:10.1073/pnas.0400054101.

41. Traag VA, Waltman L, van Eck NJ. From Louvain to Leiden: guaranteeing well-connected communities. Scientific Reports 2019 9:1. 2019;9:1–12. doi:10.1038/s41598-019-41695-z.

42. Hopcroft J, Khan O, Kulis B, Selman B. Tracking evolving communities in large linked networks. Proceedings of the National Academy of Sciences. 2004;101:5249–5253. doi:10.1073/pnas.0307750100.

43. Assessment of the control measures of the category A diseases of Animal Health Law: peste des petits ruminants. EFSA Journal. 2021;19. doi:10.2903/j.efsa.2021.6708.

44. In: Weil W, editor. Spatial Point Processes and their Applications. Berlin, Heidelberg: Springer Berlin Heidelberg; 2007. p. 1–75. Available from: 10.1007/978-3-540-38175-4_1.

45. R Core Team. R: A Language and Environment for Statistical Computing; 2021. Available from: https://www.R-project.org/.

46. RStudio Team. RStudio: Integrated Development Environment for R; 2020. Available from: http://www.rstudio.com/.

47. Cs°rdi G, Nepusz T, Traag V, Horv°t S, Zanini F, Noom D, et al. igraph: Network Analysis and Visualization in R; 2024. Available from: https://CRAN.R-project.org/package=igraph.

48. Kharchenko P, Petukhov V, Biederstedt E. leidenAlg: Implements the Leiden Algorithm via an R Interface; 2022. Available from: https://CRAN.R-project.org/package=leidenAlg.

49. Baddeley A, Turner R. spatstat: An R Package for Analyzing Spatial Point Patterns. Journal of Statistical Software. 2005;12(6):1–42. doi:10.18637/jss.v012.i06.

50. Valente TW, Coronges K, Lakon C, Costenbader E. How Correlated Are Network Centrality Measures? Connections (Toronto, Ont). 2008;28:16–26.

51. Newman M. Networks. vol. 1. Oxford University Press; 2018.

52. Candeloro L, Savini L, Conte A. A New Weighted Degree Centrality Measure: The Application in an Animal Disease Epidemic. PLOS ONE. 2016;11:e0165781. doi:10.1371/journal.pone.0165781.

53. Colman E, Holme P, Sayama H, Gershenson C. Efficient sentinel surveillance strategies for preventing epidemics on networks. PLOS Computational Biology. 2019;15:e1007517. doi:10.1371/journal.pcbi.1007517.

54. Granovetter MS. The Strength of Weak Ties. American Journal of Sociology. 1973;78:1360–1380. doi:10.1086/225469.

55. Burt RS. Structural Holes and Good Ideas. American Journal of Sociology. 2004;110:349–399. doi:10.1086/421787.

56. Hammami P, Widgren S, Grosbois V, Apolloni A, Rose N, Andraud M. Complex network analysis to understand trading partnership in French swine production. PLOS ONE. 2022;17:e0266457. doi:10.1371/journal.pone.0266457.

57. Relun A, Grosbois V, S°nchez-Vizcano JM, Alexandrov T, Feliziani F, Waret-Szkuta A, et al. Spatial and Functional Organization of Pig Trade in Different European Production Systems: Implications for Disease Prevention and Control. Frontiers in Veterinary Science. 2016;3. doi:10.3389/fvets.2016.00004.

58. Bigras-Poulin M, Barfod K, Mortensen S, Greiner M. Relationship of trade patterns of the Danish swine industry animal movements network to potential disease spread. Preventive Veterinary Medicine. 2007;80:143–165. doi:10.1016/j.prevetmed.2007.02.004.

59. Watts DJ, Strogatz SH. Collective dynamics of Ä `osmall-worldÄo networks. Nature. 1998;393(6684):440–442. doi:10.1038/30918.

60. Kim H, Anderson R. Temporal node centrality in complex networks. Physical Review E. 2012;85:026107. doi:10.1103/PhysRevE.85.026107.

61. Oettershagen L, Mutzel P, Kriege NM. Temporal Walk Centrality: Ranking Nodes in Evolving Networks. ACM; 2022. p. 1640–1650.

